# Modeling the dynamics of mouse iron body distribution: hepcidin is necessary but not sufficient

**DOI:** 10.1101/062901

**Authors:** Jignesh H. Parmar, Grey Davis, Hope Shevchuk, Pedro Mendes

## Abstract

**Background:** Iron is an essential element of most living organisms but is a dangerous substance when poorly liganded in solution. The hormone hepcidin regulates the export of iron from tissues to the plasma contributing to iron homeostasis and also restricting its availability to infectious agents. Disruption of iron regulation in mammals leads to disorders such as anemia and hemochromatosis, and contributes to the etiology of several other diseases such as cancer and neurodegenerative diseases. Here we test the hypothesis that hepcidin alone is able to regulate iron distribution in different dietary regimes in the mouse using a computational model of iron distribution calibrated with radioiron tracer data.

**Results:** A model was developed and calibrated to the data from adequate iron diet, which was able to simulate the iron distribution under a low iron diet. However simulation of high iron diet shows considerable deviations from the experimental data. Namely the model predicts more iron in red blood cells and less iron in the liver than what was observed in experiments.

**Conclusions:** These results suggest that hepcidin alone is not sufficient to regulate iron homeostasis in high iron conditions and that other factors are important. The model was able to simulate anemia when hepcidin was increased but was unable to simulate hemochromatosis when hepcidin was suppressed, suggesting that in high iron conditions additional regulatory interactions are important.

## Background

Iron deficiency affects 24.8 % of the population worldwide [1] while hereditary hemochromatosis, an iron overload disease, affects 1 in 200 whites of northern European ancestry [2]. Iron is an essential element, playing a role in oxygen binding and electron transport. Despite its importance, iron can be dangerous when free or poorly liganded in solution as it can catalyze the production of hydroxyl radical from hydrogen peroxide. Indeed there is strong evidence implicating iron together with reactive oxygen species in a wide range of diseases, including cancer, diabetes, and neurodegenerative diseases [3]. Mechanisms have evolved to maintain iron homeostasis and store iron in non-toxic forms. Because pathogens grow faster than mammalian cells, mechanisms have also evolved that sequester iron from circulation during infections [4, 5], limiting proliferation of the infectious agent. Given its simultaneous essentiality and toxicity, it is not surprising that iron homeostasis requires efficient control mechanisms. These mechanisms must be able to damp the effects of deficient or excessive iron intake in normal individuals. Improper regulation of iron homeostasis leads to iron-deficiency anemia, or to hemochromatosis due to iron excess.

The current understanding of iron regulation is by control of its absorption and release from organs such as the liver and spleen [6]. This is because there is no known active route for excretion of iron in mammals. The hormone hepcidin (HAMP), synthesized and secreted mainly by the liver [7], is believed to be the main regulator of iron [5, 6]. The purpose of the present work is to quantitatively test the regulation of iron distribution by hepcidin through the development of a computational dynamic model of iron distribution in the mouse. The strategy is to calibrate the model with data from mice eating an adequate iron diet, and then test its performance against data of mice eating deficient and rich iron diets. The results of this model will confirm or refute the hypothesis that hepcidin alone can explain the distribution of iron at different dietary levels. The plasma iron level depends on intestinal absorption of iron, red blood cell recycling and import and export from various organs, particularly the liver and spleen. Iron excretion occurs only through sloughing of enterocytes, skin and hair shedding, sweating, and blood loss. Dietary iron appears mainly in the form of Fe^3+^ and it is reduced to Fe^2+^ by duodenal cytochrome-b (DCYTB) and other brush border ferrireductases. Fe^2+^ is then imported into enterocytes by the divalent metal transporter 1 (DMT1). In enterocytes it is exported to the serum through ferroportin (FPN, or SLC40A1), the only known iron exporter in mammalian cells [8]. In the plasma, iron binds to the transport protein transferrin (TF) which donates iron to various cell types through the transferrin receptors (TFR1, TFR2). Iron can also be exported from hepatocytes and macrophages into the plasma through ferroportin. Excess iron in the body is stored mainly in the liver, in the form of ferritin.

Hepcidin induction increases with high serum iron levels [9, 10] and decreases with iron deficiency [10, 11]. Hepcidin promotes the degradation of ferroportin [12], thus higher levels of hepcidin result in lower levels of ferroportin, with the consequent accumulation of iron in tissues and lower rate of dietary iron acquisition, and a decrease in plasma iron level. This effect of iron immobilization is important also in immune defense to limit microbial pathogen growth and happens due to the induction of hepcidin by cytokines. The current understanding of the role of hepcidin in regulating iron homeostasis is therefore that a) at high iron loads hepcidin is induced thus decreasing absorption of iron (reduction of enteric ferroportin), b) at low iron loads hepcidin is reduced, thus allowing transfer of iron from the diet to the plasma. Additionally, poor regulation of hepcidin results in disease: c) chronically low hepcidin or defects in upstream signaling elements (such as HFE, TFR2, HJV genes) causes iron overload typical of hemochromatosis, and d) chronically high hepcidin (e.g. induced by inflammatory cytokines) causes anemia.

The conceptual model of iron regulation by hepcidin described above is widely illustrated in various reviews [5,6,12–14]. While conceptual models are useful to reason about regulation, ultimately they must be tested quantitatively. This is best achieved by mathematical models and computer simulation. Several computational models addressed particular aspects of iron biochemistry, such as iron loading of ferritin [15], erythropoiesis [16–18], or liver hepcidin regulation [19] but few consider iron distribution across the body. A good systemic iron model should be calibrated with physiological data, reflect the dynamics of iron in response to dietary and genetic perturbations, and be validated with independent experiments. Lao and Kamei [20] developed a compartmental model of systemic iron, however it was not calibrated or validated with experimental data. Quantification of iron distribution across the body requires data from radiolabeled iron uptake in different organs [21]. Lopes *et al.* [22] built three steady-state flux distribution models based on a data set of radioactive iron distribution in plasma and 15 other organs in mice under adequate, iron-deficient, and iron-rich diets [23]; each model describes the steady state exchange fluxes of iron between organs in one of the three dietary regimes [22]. However these models are unable to test the hepcidin regulation hypothesis as they do not include this hormone. They are also not able to simulate transitions between diet regimes, as each one has a distinct set of parameter values (first order rate constants), fitted independently from different time courses of radioactive iron distribution, one for each dietary regime.

The dynamic model of systemic iron presented here includes the action of hepcidin, as currently understood, and is calibrated against the Schümann *et al.* dataset [23]. In order to increase the molecular detail of the current model relative to those of Lopes *et al.* [22], we included known or hypothesized details of iron metabolism and further relevant data collected from other studies. The strategy taken here was to estimate parameters (calibrate the model) against the adequate iron diet dataset and then to investigate if the same model would be able to simulate the data for iron-deficient and iron-rich diets, by allowing only changes in two parameters: dietary iron loading and hepcidin synthesis rate. If the model is able to reproduce the deficient and rich diets in such a way then this shows that the conceptual model of iron regulation by hepcidin is possibly correct (or at least compatible with the data). The model can be further tested/validated by simulating the causes of chronic anemia and hemochromatosis and checking whether the results are consistent with the corresponding phenotypes.

## Methods

### Modeling framework and computations

The model is based on ordinary differential equations (ODE), with each pool (species) being modeled by one ODE composed of positive terms that account for the production of the species and negative terms that account for the consumption of the species. Model creation and computations were carried out with the software COPASI version 4.16 (http://copasi.org) [24] running on computers with Apple OS X, Microsoft Windows and Linux operating systems. Cell Designer [25] and Inkscape were used to draw SBGN [26] diagrams. Parameter estimation was carried out to fit the data published by Schümann *et al.* [23] by least-squares minimization using various optimization algorithms (see Table 2) in the COPASI parameter estimation framework. The present model is provided as supplementary data to this article in SBML [27] and COPASI formats, in two versions: one including the radioactive tracer species and another only with non-radioactive species. Two models for iron-deficient and iron-rich diets are also provided. These models were deposited in BioModels (http://www.ebi.ac.uk/biomodels) [28] and assigned identifiers MODEL1605030002, MODEL1605030003, MODEL1605030004, MODEL1605030005.

**Table 2.**
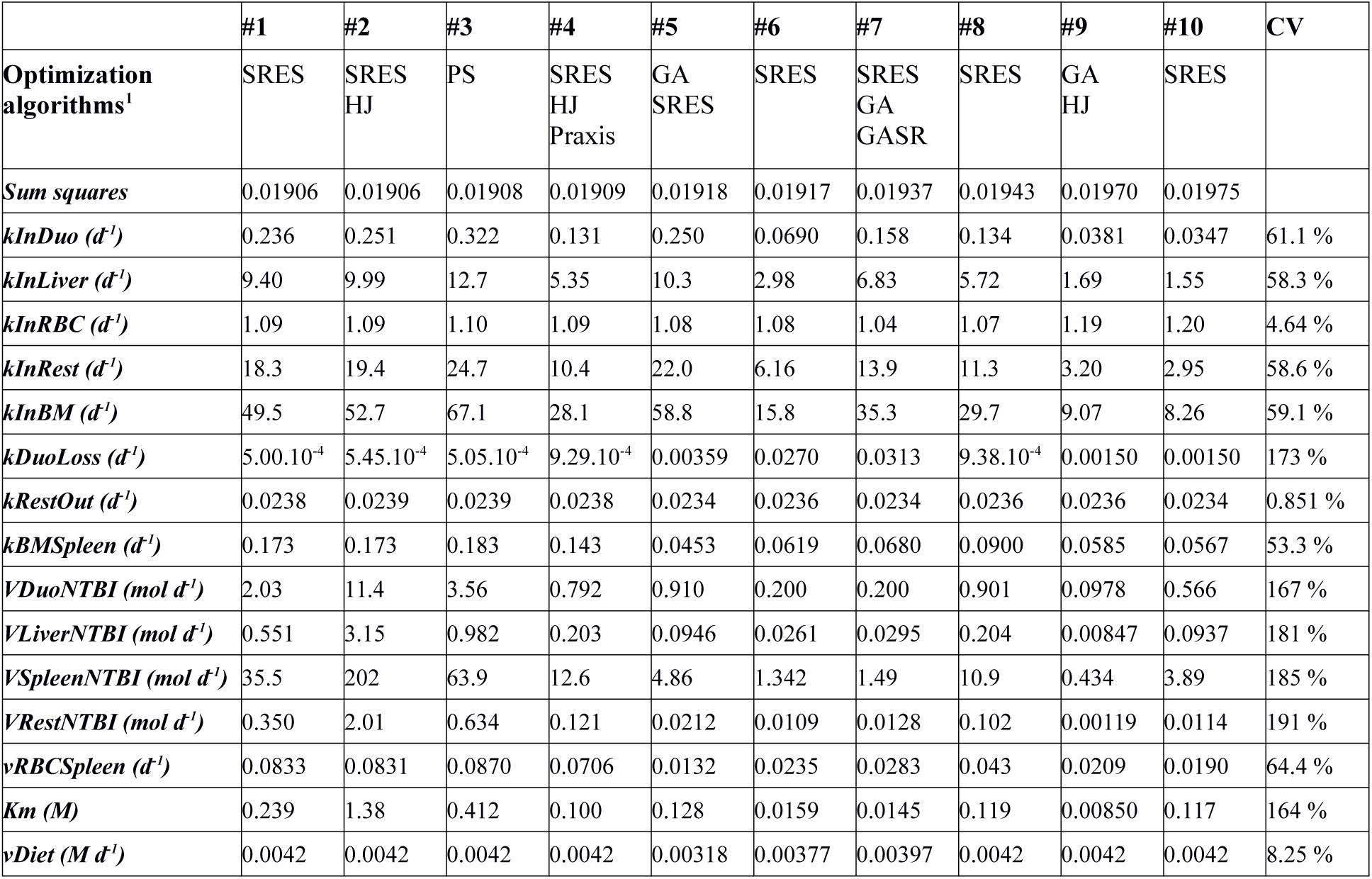
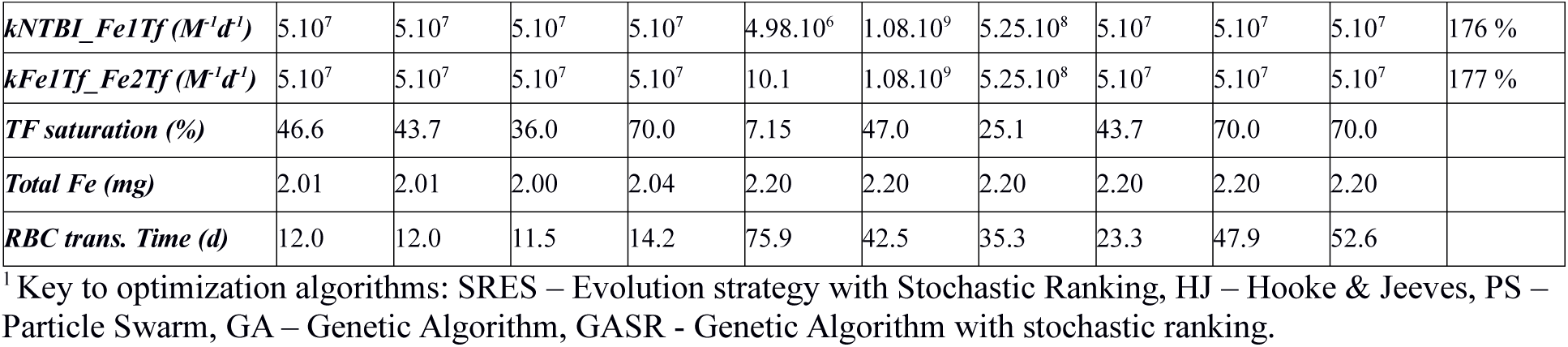
Independent parameter estimates and properties of their models. Model #6 was chosen due to its RBC transition time and Tf saturation being closer to known values in mice. Coefficient of variation (CV) is expressed for each of the estimated parameters across all independent estimates.

### Model and data

The intention with this model is to represent the iron homeostasis in the mouse in the presence of adequate iron diet. The data from Schümann *et al.* [23] measures the decay of a small single injection of radioactive iron (^59^Fe); the dose is sufficiently small for the perturbation in total iron to be negligible, thus the rates that were estimated from these data in [22] correspond to steady state fluxes of iron between compartments. While the models of Lopes *et al.* [22] only account for the radioactive iron, the present model accounts for both radioactive *and* non-radioactive iron. This was achieved by duplicating all reactions in the model, one for the radioactive form and another for the non-radioactive form, and making sure that both share the same value of rate constants (*i.e.* assuming that kinetic isotope effects in the system can be ignored [29]). Where necessary the competitive effects between the two iron pools are taken into account.

### Model compartments

While Schümann *et al.* [23] provide data for plasma and 15 different organs, in the present model we sought some simplification by only considering the most important organs for iron homeostasis. Thus the present model represents: plasma, liver, duodenum, bone marrow, red blood cells, spleen and a compartment for the rest of the body that accounts for the aggregate of iron in stomach, intestine, integument, muscles, heart, fat, lungs, kidneys, brain and testes (*i.e.* the ^59^Fe content in these compartments in Schümann *et al.* [23] were here added into one single ^59^Fe species). As the main routes of iron excretion from the body are enterocyte sloughing, skin desquamation, and hair loss, we reflect these facts by iron export from the duodenum and rest of body to an additional compartment “outside”, which is only important for parameter estimation. Fig. 1 represents the model compartments and the movement of iron between them.

**Figure 1.**
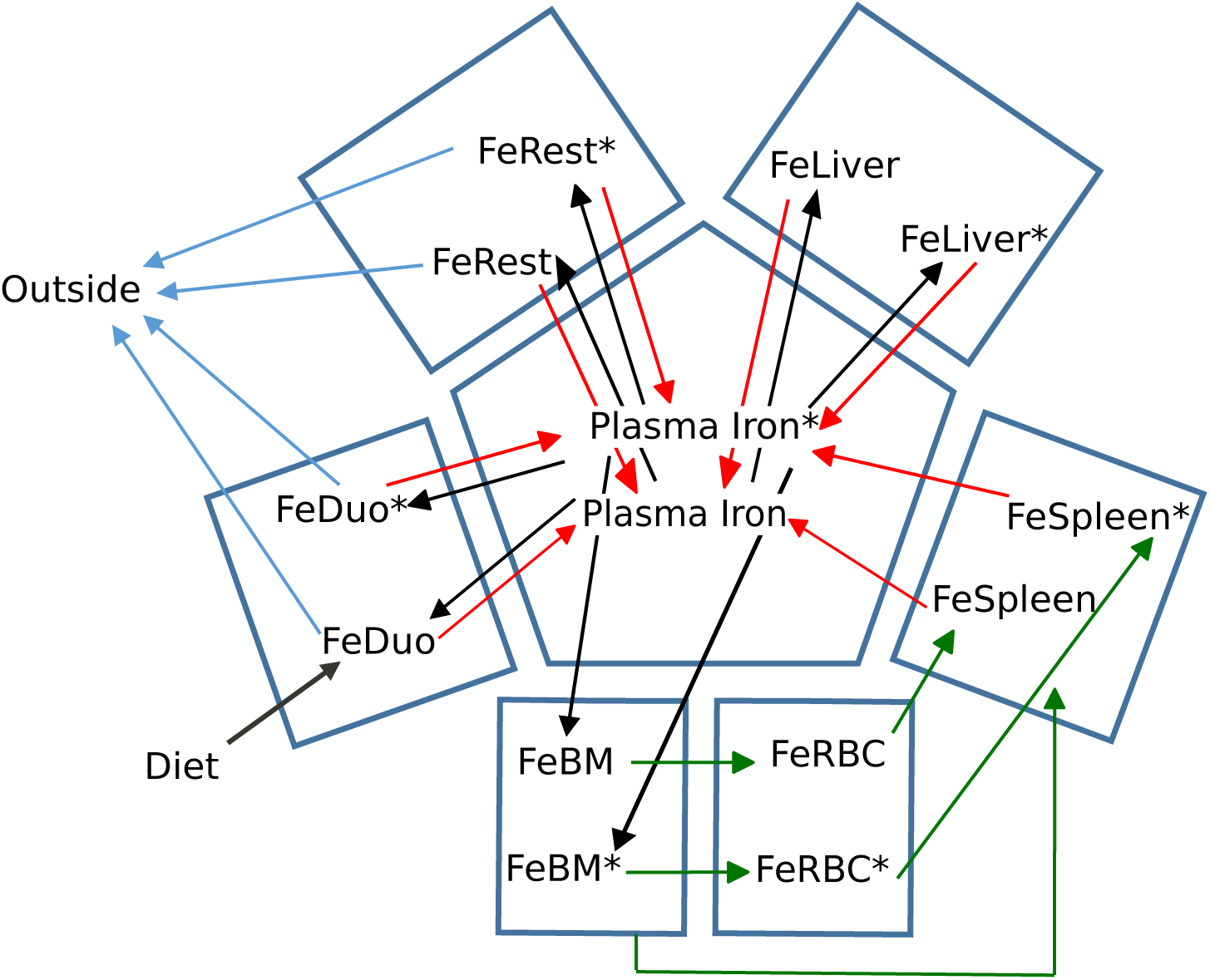
Movement of iron between compartments in the model. Arrows indicate the flow of iron. The symbol ‘*’ marks the radioactive species. Black arrows represent transfer of iron from transferrin to tissues, blue arrows represent transfer of iron from tissues to plasma through ferroportin, green arrows represent erythroid cell dynamics, and grey arrows represent loss of iron.

Since we desired to calibrate the model with absolute concentrations, it was required to estimate volumes for each of these organs considered. We converted the masses of organs provided in [23] to volumes using densities of equivalent human organs, as published by the International Commission on Radiological Protection [30]. Ideally this conversion should have used mouse organ densities, but these could not be found in the literature (this error is expected to be small). The volumes considered are listed in Table 1.

**Table 1.** Mass, density and volume of model compartments.

| Compartment | Mass (g) <sup>1</sup> | Density (g.cm<sup>-3</sup>) <sup>2</sup> | Volume (l) |
| --- | --- | --- | --- |
| Bone Marrow | 0.21 | 0.98 | $2.1 \cdot 10^{-4}$ |
| Duodenum | 0.04 | 1.04 | $3.9 \cdot 10^{-5}$ |
| Liver | 1.22 | 1.05 | $1.2 \cdot 10^{-3}$ |
| Plasma | 1.36 | 1.03 | $1.3 \cdot 10^{-3}$ |
| RBC | 0.71 | 0.90 | $7.9 \cdot 10^{-4}$ |
| Rest of body<sup>3</sup> | | | $2.0 \cdot 10^{-2}$ |
| Spleen | 0.07 | 1.04 | $6.7 \cdot 10^{-5}$ |

<sup>1</sup>From Ref. [23].
<sup>2</sup>From Ref. [30].
<sup>3</sup>Volume was calculated as the sum of volumes of all organs considered in [23] (calculated using the respective densities from [30]) minus the volumes of the other rows in this table.

### Chemical species

With the exception of plasma, each model compartment includes one species representing that organ′s total non-radioactive iron content and another one representing its total radioactive iron content. Using the symbol ′*′ to denote the radioactive forms, there are FeBM and FeBM* in the bone marrow, FeDuo and FeDuo* in the duodenum, FeLiver and FeLiver* in the liver, FeSpleen and FeSpleen* in the spleen, FeRBC and FeRBC* in the red blood cells, FeRest and FeRest* in the rest of the body, and FeOutside* that accounts for the excreted iron. In the plasma compartment the model represents several additional chemical species: hepcidin, apo-transferrin (Tf), non-transferrin bound iron (NTBI) and its radioactive equivalent (NTBI*), transferrin with a single non-radioactive bound iron ion (Fe1Tf), and with a single radioactive iron bound ion (Fe1Tf*), transferrin with two non-radioactive bound iron ions (Fe2Tf), with two radioactive bound iron ions (Fe2Tf**) and with one radioactive and one non-radioactive bound iron ions (Fe2Tf*). The reactions between these plasma species follow mass action kinetics as they represent simple binding and dissociation.

### Reactions and reaction kinetics

Fig. S1 depicts the full set of reactions of the model using the SBGN standard [26]. The number of reactions makes the diagram somewhat difficult to understand, thus Fig. S2 illustrates the model without considering the radioactive species (which were only needed for parameter estimation).

#### Transferrin and NTBI reactions

Binding of NTBI or NTBI* to Tf happens through reversible mass action reactions. By considering all the explicit mass action reactions between all radioactive and non-radioactive species, the mutual competitive effect of NTBI versus NTBI* is captured naturally. Given that the *K*_*d*_ for iron binding to transferrin is in the order of 10^−22^ M [31] we considered these reactions irreversible to keep the model simpler.

#### Transferrin-mediated iron import to organs

The model considers that iron import from the plasma into the bone marrow, liver, duodenum and rest of body happens only via the trasferrin-bound iron species, releasing the iron ions into these organs while apo-transferrin (Tf) remains in the plasma. These organs may receive one iron ion (from Fe1Tf or Fe1Tf*) or two (from Fe2Tf, Fe2Tf* or Fe2Tf**). These iron import reactions are considered as irreversible mass action in the model. Arguably these processes are saturable given that they are mediated by receptors (TfR1 and TfR2); however since the model does not consider their regulation they were left as first order reactions, assuming that there is always enough capacity of transferrin receptors for the supply of iron from transferrin

#### Ferroportin-mediated iron export

Iron is exported to the plasma through ferroportin, which is the only known transporter capable of exporting iron from mammalian cells [8]. Ferroportin iron export is represented by saturable kinetics (Equation 1); the rate law includes a term for competitive inhibition to represent the effect of the radioactive and non-radioactive species competing for the same transporter (obviously a symmetric effect, *i.e.* the *K*_*m*_ for the substrate is the same as the *K*_*i*_ for the competitive inhibitor). The rate law also contains a non-competitive inhibition term representing the effect of hepcidin. The kinetic effect of non-competitive inhibition is equivalent to reducing the apparent *V*_*max*_ and this reflects the mode of action of hepcidin since it promotes removal of ferroportin molecules (the enzyme of this reaction), thus causing a lowering of the apparent *V*_*max*_. The model includes ferroportin export reactions to the plasma from the duodenum, spleen, liver and rest of body. The rate of the ferroportin-dependent iron export is thus:

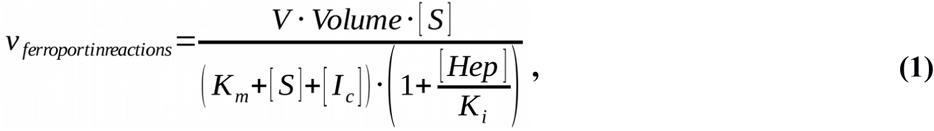

where *V* is the limiting rate (*V*_*max*_, proportional to the concentration of ferroportin), *Volume* is the volume of the compartment where the substrate and competitive inhibitor are located, [*S*] is the substrate concentration (*e.g.* FeSpleen), *K*_*m*_ is the Michaelian constant of the substrate (also the competitive inhibition constant of the competitive inhibitor), [*I*_*c*_] is the competitive inhibitor concentration (*e.g.* FeSpleen*), [*Hep*] is hepcidin concentration and *K*_*i*_ the hepcidin non-competitive inhibition constant. The value of the *K*_*m*_ and *K*_*i*_ constants is the same for all the ferroportin reactions since these parameters reflect properties of the protein. On the other hand the parameter *V* is estimated independently for each of the four organs, since it reflects the ferroportin protein level in each one and therefore is a characteristic of the tissue. Note that the rate of reaction is expressed in extensive units (amount per time) rather than intensive units (concentration per time), this is because this reaction crosses two compartments of different volume. COPASI forces such reactions to be expressed in extensive units in order to properly take into account the effect of the different volumes (in fact COPASI internally calculates all ODEs in units of amount per time).

#### Erythroid cell dynamics

About 20-25% of the total body iron cycles between the plasma, bone marrow, RBC and spleen. Iron is acquired by bone marrow cells that eventually maturate into RBC. This represents an iron transfer from one compartment to the other which is modeled here as a first order reaction for both radioactive and non-radioactive iron (same value for their rate constants). Similarly red blood cells are eventually destroyed by spleen macrophages that keep their iron. This process is also reflected in the model by an irreversible mass action reaction from the RBC iron species to spleen iron species (again both radioactive and non-radioactive with same rate constants). Spleen macrophages are also known to destroy some pre-matured erythroid cells [32] so the model includes a direct iron uptake from the bone marrow to the spleen (as in [22]), also represented by irreversible mass action kinetics. There is no acquisition of iron from plasma by the RBC; all their iron was already acquired in the bone marrow.

#### Dietary iron import

The entry of dietary iron into the circulation happens through the duodenum. The entry of iron into the enterocytes of the duodenum is here represented by a constant flux reaction that represents the time average rate of import of dietary iron rather than simulating specific meals as discrete time events; given that the amount of iron absorbed per meal is a small amount compared to the total iron in the system, such discrete events would have a small effect and in fact are not detectable in the experimental data used here – thus this simplification is not expected to have significant effect in the results. Considering the import of iron as a continuous process allows this model to define a steady state for which there are many useful analysis tools (the same assumption is also included in [22]). In other uses of the model, this reaction can be set to any other desired flux to represent iron-deficient or iron-rich diets. It could also be switched on and off at discrete time points to simulate individual meals.

#### Iron loss from the body

Because the enterocytes in the duodenum have a short half-life there is a considerable loss of iron through their sloughing, which is therefore represented as an irreversible mass action reaction from this compartment to the “outside” compartment. A similar process happens to skin and hair cells, which eventually are shed from the body and thus reduce the total iron. Therefore the model also contains an irreversible mass action reaction from the rest of body to the “outside” compartment (both of these reactions are, of course, mirrored by equivalent ones with the radioactive form of iron).

#### Hepcidin

The model considers hepcidin to be synthesized at a constant rate and degraded by a first order reaction. The values of the parameters of these reactions were set to result in a level of hepcidin in the nM range. The rate of synthesis is then adjusted to fit the data in the deficient and rich iron diets. Ideally a function would reflect the level of hepcidin given the level of transferrin-bound iron, however we do not know of data that allows proper calibration of such a function as the only data available for mouse hepcidin consists of mRNA levels in tissues, rather than the level of circulating active hormone peptide.

#### Initial conditions

The experiments are carried out by injecting a certain small amount of radioactive iron to male young adult mice that have been kept for 35 days in a certain iron diet (adequate, deficient or rich). Thus one expects this perturbation to occur in a quasi steady-state of iron and transferrin in the body. We assumed a total concentration of transferrin of 3.87 × 10^−5^ M [33] and initialized this entirely in the apo-transferrin form. Since there is conservation of transferrin in the stoichiometry of this model, this ensures the total Tf to remain at this value. The total iron amount in normal mice is known to be around 2 *mg* [34], but since there is no conservation of iron mass in this model, we have no direct control over this value – therefore we set the following constraint on the parameter estimation: 1.8 *mg* < *TotalFe* < 2.2 *mg* (where *TotalFe* is the total iron amount obtained by summing all variables that reflect iron concentration). This results in a model that fits the data while also having a total amount of body iron within 10% of the expected value.

Because we do not know the amounts of non-radioactive iron nor the partition of transferrin in its various iron-bound forms in the initial state, these had to be estimated from the data together with the rate constant values. To achieve that, the time course was first set to take a long “lead-in” time (5000 days) before the radioactive iron is injected. Since the parameter estimation starts only when the radioactive iron is injected, at that point the various non-radioactive species are in a quasi steady-state. Once a satisfactory fit to the radioactive data is obtained we then repeated the process by setting the initial concentrations of all species to be what they were in the quasi steady-state obtained previously, but this time only having a lead in time of 35 days (equivalent to the experiment lead in time for each diet).

## Results

### Parameter estimation for adequate iron diet

The values of kinetic constants were estimated in order to match the ^59^Fe time-course data in the iron-adequate diet experiment. As reported above, a constraint was added to keep the total body iron at 2 ± 0.2 *mg*. To reduce parameter unidentifiability, we set the rate of iron import into the body (*vDiet* parameter, the rate of the transfer of iron from the outside into the duodenum) to the value 0.0042 mol day^-1^ which is in the range of expected iron turnover in mice [35].

Several independent parameter estimates were carried out using different optimization algorithms or combinations thereof. A list of all the parameter estimates obtained is included in Table 2. The optimization algorithm that was most successful for this problem was the SRES algorithm [36], most likely due to the use of a nonlinear constraint [37]. From all the solutions, we picked the one that has a transition time [38, 39] for the iron in red blood cells close to the known average lifetime of these cells in mice (40 days [40–42], the chosen parameter set results in a transition time of 42.5 days). Parameter set #6 in Table 2 is therefore the one used hereafter, and used in the models included in supplementary data (MODEL1605030002 and MODEL1605030003 in BioModels).

An important question about this parameter estimation is whether the parameters are identifiable. From Table 2 we can see that a couple of parameters have similar estimated values across independent runs (small coefficient of variation, CV): the rate constant for iron loss from the rest of body compartment (*kRestOut*), the rate of maturation of erythrocites (*kInRBC*) – thus we have high confidence on these values. On the other hand there is likely a considerable degree of dependence between the other parameters, yet the conclusions of this study are robust to this parameter variation, as shown in supplementary Figure S3 depicting simulations by all 10 models of Table 2.

Figure 2 shows that the model results are in good agreement with the experimental time-courses of ^59^Fe in the various tissues compartments. The model largely reproduces the observations from mice: radioactive iron in the plasma drops sharply as it rapidly transported to other compartments. The liver and the rest of body compartments show marked increase in ^59^Fe level followed by a gradual decline as the iron is exported back into the plasma through ferroportin. Bone marrow rapidly takes up to 60% of the injected ^59^Fe which then declines as it is utilized in the RBC production, reaching a level of 60% in a few days. However, if simulating for a longer time, say 300 days, the model shows a gradual disappearance of ^59^Fe as it is excreted from the body supporting a similar experimental observation by Stevens *et al.* [43]. In three compartments the model does not fit the data perfectly: the simulated spleen ^59^Fe has the initial overshoot smaller than that displayed by the data; in the rest of the body the simulation overestimates the initial amount of ^59^Fe; in the bone marrow the simulation predicts too fast a decay of ^59^Fe, which in the data takes longer. But these deviations are minor and in the case of the spleen the simulation is within one standard deviation of the data.

**Figure 2.**
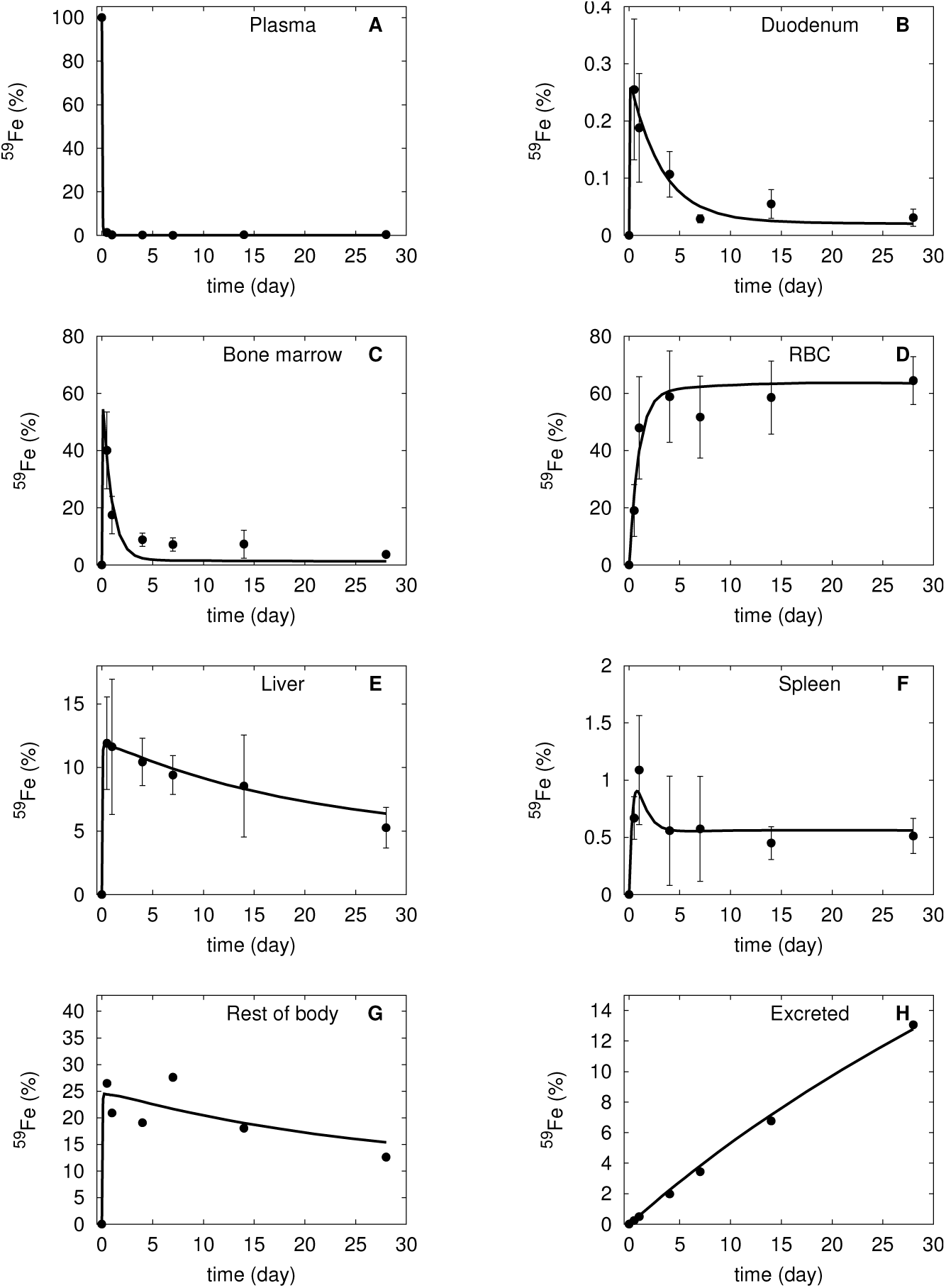
Simulation of the calibrated model (#6 in Table 2) against the experimental data of ^59^Fe injection in mice under adequate iron diet. Continuous line is the model simulation, while filled circles represent the data from [23] (vertical bars represent one standard deviation). The abscissa represents the proportion of total injected ^59^Fe, the ordinate is time after injection. **A** plasma, **B** duodenum, **C** bone marrow, **D** red blood cells, **E** liver, **F** spleen, **G** rest of body, and **H** excreted. Note that the standard deviation in panel **A** is smaller than the size of the symbols, while in panels **G** and **H** it was not indicated as it is very large because these data are algebraic sums of various terms.

Overall the model is quite capable of reproducing the dynamics of radiolabeled iron in the experiment. The parameter values for the rate constants are therefore adopted as the model for mice under an adequate iron diet. The model is supplied in its entirety in the COPASI file Parmar2016-AdequateTracer.cps and without the tracer species in the file Parmar2016- AdequateNoTracer.cps (and corresponding SBML versions of these files).

### Iron-deficient diet

In accordance with the experimental protocol, the model for iron-adequate diet is simulated with iron-deficient diet (reduced *vDiet*) for 5 weeks before simulating the pulse of ^59^Fe. First we show that changing the *vDiet* parameter alone does not successfully reproduce the observations from iron-reduced diet mice. The simulated iron level is too high in the liver, rest of body, and spleen and too low in the RBC, compared to the corresponding experimental values (see Figure 3, dotted lines). Then we also allowed the rate of synthesis of hepcidin to be adjusted in order to better fit the data since this affects its steady state level and therefore the rate of ferroportin-dependent iron export (since hepcidin is a non-competitive inhibitor of these reactions). Once the hepcidin is also reduced the simulation then better matches the iron levels in the liver and rest of body. Nevertheless, the amount of simulated iron in the spleen is still not a good match to the experimental data.

**Figure 3.**
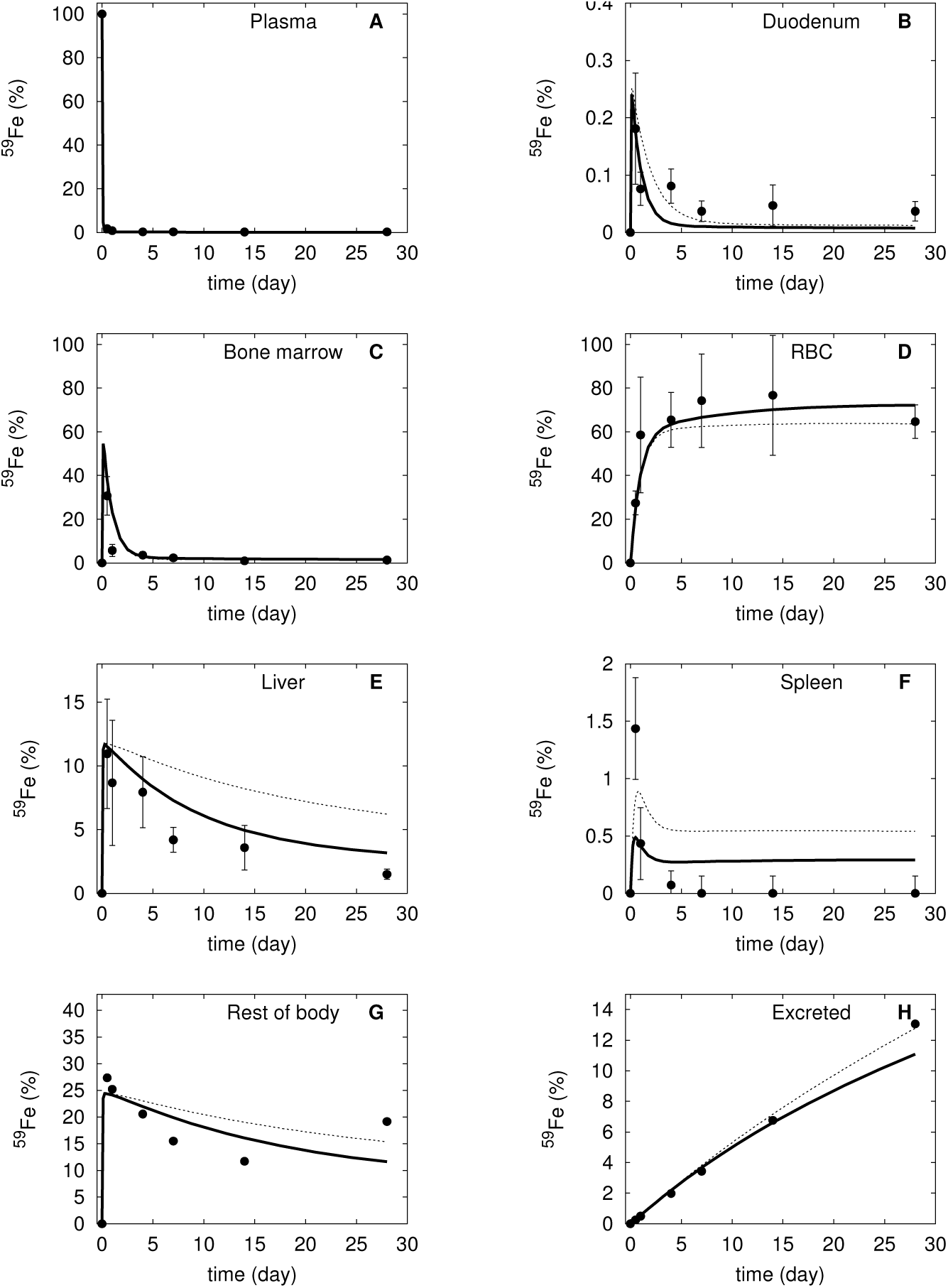
Simulation of the calibrated model (#6 in Table 2) against the experimental data of ^59^Fe injection in mice under iron-deficient diet. The dotted line represents the simulation when only the diet was adjusted, the bold line is the model simulation when both diet and hepcidin synthesis rate were adjusted. Filled circles represent the data from [23] (vertical bars represent one standard deviation). The abscissa represents the proportion of total injected ^59^Fe, the ordinate is time after injection. **A** plasma, **B** duodenum, **C** bone marrow, **D** red blood cells, **E** liver, **F** spleen, **G** rest of body, and **H** excreted. Note that the standard deviation in panel **A** is smaller than the size of the symbols, while in panels **G** and **H** it was not indicated as it is very large because these data are algebraic sums of various terms.

The deficient diet model, in its version without tracer, was then used to simulate the outcome after 100 days under this diet (Table 3). The result is that the total iron mass in the body would reduce by 23% (from 2.2 to 1.7 mg) while the transferrin saturation would decrease from 47% to 37%. The iron excretion rate was of 5 µg/day under this diet, which is in general agreement with data from [34] which reported an excretion rate of 8 µg/day under a milk diet. It also underlines the fact that since hepcidin can only limit the entry of iron to the plasma, an iron deficient diet will inevitably lead to loss of iron.

**Table 3.** Characteristics of the three simulated diets. The table lists the simulated values of various physiological variables after 100 days in the specified diet.

| Variable | Adequate diet | Deficient diet | Rich diet |
| --- | --- | --- | --- |
| Tf saturation | 47 % | 37 % | 44 % |
| Total iron | 2.2 mg | 1.7 mg | 2.2 mg |
| [Hepcidin] | 23 nM | 11 nM | 31 nM |
| [NTBI] | 33 nM | 22 nM | 30 nM |

### Iron-rich diet

Simulation of the iron rich diet experiment follows a similar approach as the deficient diet. We set the different diet regime 5 weeks before the radioactive tracer pulse is applied. First we attempted to fit the data only by changing the rate of iron entry in the duodenum (*vDiet* parameter). In this case the fit is also not accurate enough (Figure 4), with simulated RBC iron tracer much higher than actual measurements, while the simulated liver iron tracer is much lower than measurements.

**Figure 4.**
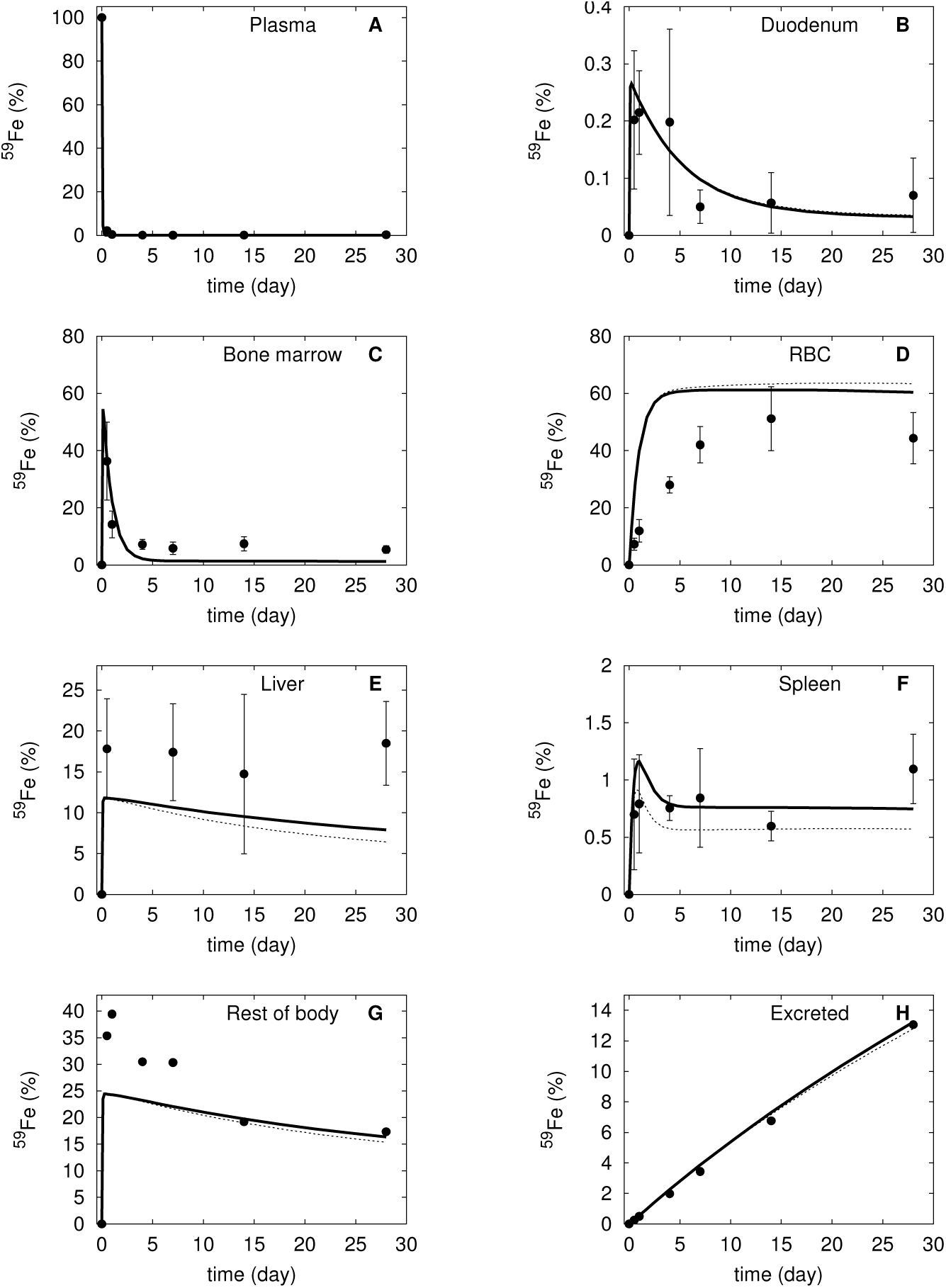
Simulation of the calibrated model (#6 in Table 2) against the experimental data of ^59^Fe injection in mice under iron-rich diet. The dotted line represents the simulation when only the diet was adjusted, the bold line is the model simulation when both diet and hepcidin synthesis rate were adjusted. Filled circles represent the data from [23] (vertical bars represent one standard deviation). The abscissa represents the proportion of total injected ^59^Fe, the ordinate is time after injection. **A** plasma, **B** duodenum, **C** bone marrow, **D** red blood cells, **E** liver, **F** spleen, **G** rest of body, and **H** excreted. Note that the standard deviation in panel **A** is smaller than the size of the symbols, while in panels **G** and **H** it was not indicated as it is very large because these data are algebraic sums of various terms.

Then we proceeded to allow the hepcidin synthesis rate to vary as well. In this case the overall fit improved only by a minimal amount. Essentially the simulation is unable to fit the measurements as it accumulates too much iron in the RBC and not enough in the liver. Figure 4 shows that the action of hepcidin in the model, even though in the right direction, is not enough to explain the data. Note that the rate of synthesis of hepcidin was not limited in the fit, and the value obtained is the one that gives the best overall fit to all the curves (i.e. a higher rate of hepcidin synthesis resulted in worse fits).

When using the last model (with altered *vDiet* and hepcidin synthesis rate) to simulate the effect of this diet regime for 100 days, we observe that despite an increase in hepcidin by 35% (from 23 to 31 nM), the total iron, transferrin saturation and NTBI concentration stayed essentially unchanged (Table 3). These results are also *not* in line with previous observations.

At this point it is important to investigate whether other parameter sets from Table 2 would better fit the iron-rich diet observations than #6. Thus we further tested the predictions of the other parameter sets from Table 2 for the iron-deficient diet data. However, as shown in Fig. S3 (supplementary data), no parameter sets were able to better match these data than parameter set #6 discussed above. Another possibility could be that using all three data sets (adequate, iron-deficient and iron-rich diets) for parameter estimation would result in a model that would explain all of the data well. We then tried to estimate parameters by fitting all data sets simultaneously (only allowing *vDiet* and the synthesis rate of hepcidin to be different for each diet). But even in this case the best fit model still shows an inability to match the observations for the iron-rich diet, particularly for the RBC and liver iron contents (Figs S4-S6 in supplementary data). This strengthens the conclusion that a model where only changes in hepcidin synthesis rate regulate iron distribution is not sufficient to explain the rich diet observations – a conclusion that is independent from the exact parameter estimates as all independent models led to the same result.

Finally we decided to take an alternative approach, by fitting the model to the iron-rich diet only and validating it to the adequate and iron-deficient diets. Table S1 (supplementary data) depicts a set of models that fit the iron-rich data, but as shown in Figs S8 and S9 these models cannot explain the other two data sets. A comparison of parameter values between the models calibrated against the iron-rich diet (Table S1) and those calibrated against the adequate iron diet (Table 2) was carried out *via* a pair-wise *t*-test (Table S2). The most significantly different parameters between the adequate and rich diets were *kRestOut* (the excretion rate through the rest of body), *kInRBC* (the intake rate of iron by the RBC) and *kInliver* (the intake rate of iron by the liver). This suggests that additional regulation on these processes may be taking place, alongside with the hepcidin regulation, under iron rich conditions.

### Simulating the anemia of chronic disease

Anemia of chronic disease is caused by elevated levels of hepcidin, which immobilize iron in tissues and prevents transfer of enterocyte iron to the plasma. This disease can be simulated using our adequate iron model by increasing the rate of synthesis of hepcidin. We manipulated the hepcidin synthesis rate in order to increase the hepcidin concentration five-fold and observed how the model evolves during 365 days, still with an adequate iron diet (Figure 5). The simulation results qualitatively match with the known physiological indicators of this disease. The plasma iron (both transferrin-bound and NTBI), bone marrow iron and RBC iron decreased by approximately 3-fold, while the iron in the rest of body decreased about 1.5-fold. On the other hand iron accumulated in the duodenum (8-fold), liver (2-fold), and spleen (2-fold). In the spleen there is a sharp increase in the time frame of 50 days, reaching 10x the initial concentration, followed by a slower relaxation. In the liver there is also an overshoot, though that dynamics is slower, with the peak at around 180 days. The total body iron decreased by 2-fold.

**Figure 5.**
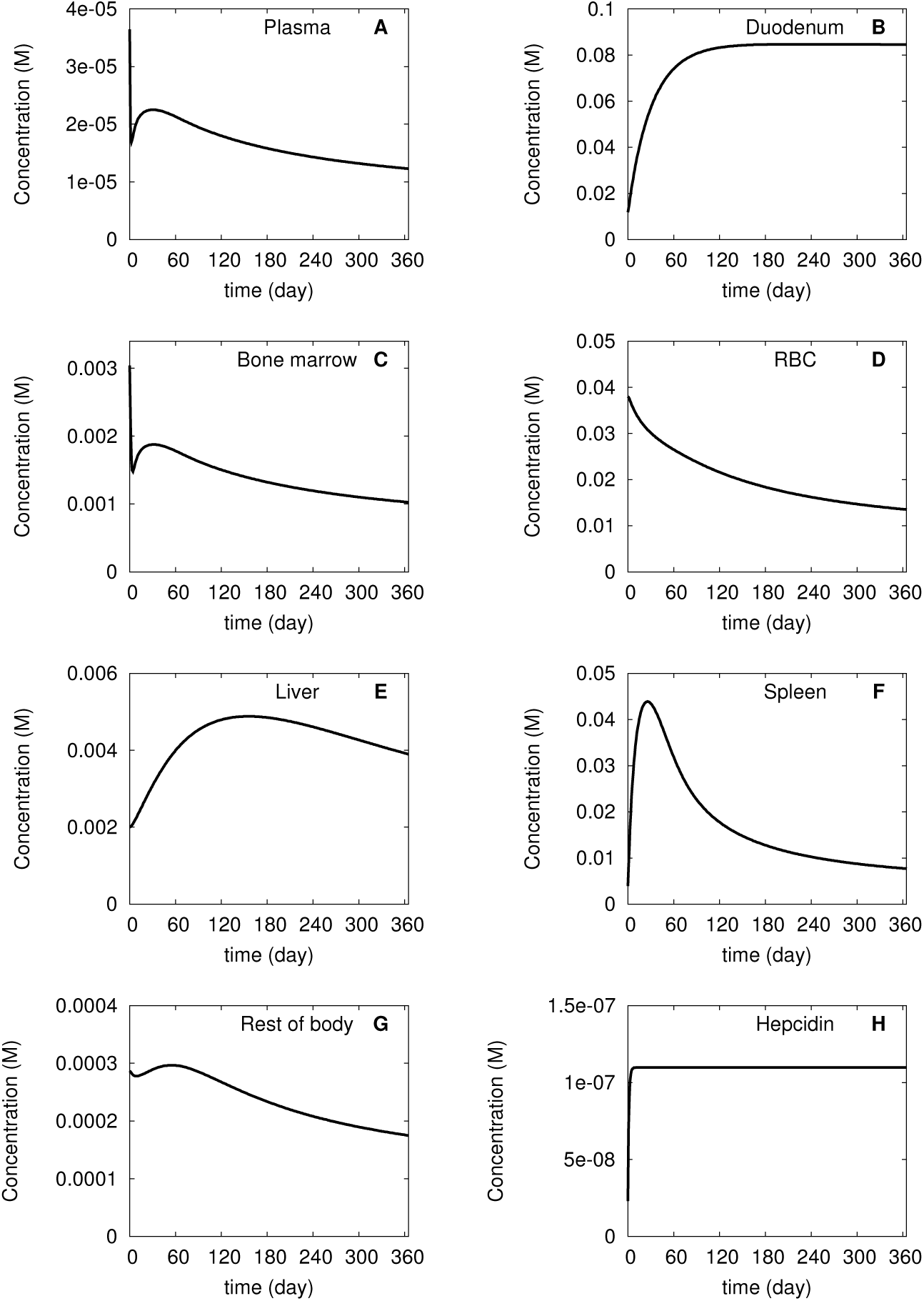
Simulation of anemia of chronic disease. The model calibrated for the adequate iron diet was used (#6 in Table 2). At time zero the hepcidin level was increased five-fold and maintained at that level for 365 days. The abscissa represents concentration and the ordinate the time. **A** plasma, **B** duodenum, **C** bone marrow, **D** red blood cells, **E** liver, **F** spleen, **G** rest of body, and **H** hepcidin.

## Discussion

Models constitute a quantitative framework for reasoning about phenomena. George Box famously said that “all models are wrong, but some are useful” [44]. Indeed the current model is, of course, wrong in many details, but it is useful in revealing the role of hepcidin in iron homeostasis and the extent to which it does *not* seem to explain all observations. The main finding of this work is that considering hepcidin as the single iron regulator does not allow us to explain the body iron distribution under a broad range of iron diets. When the model is calibrated against the adequate diet data, it overestimates the amount of iron in red blood cells and underestimates the amount of liver iron under the iron-rich diet. If the model would be calibrated to the iron-rich data, it then understimates the iron in RBC and overestimates the iron in the liver of the adequate diet. These failings of the model suggest other important factors to be at play.

Before we consider the failings of the model, it is worth pointing out that it is successful in explaining the iron distribution under adequate iron diet, and its prediction for iron-deficient diets is not too far from reality. This allowed us to attempt to simulate anemia of chronic disease, a condition that arises from abnormally elevated hepcidin, usually caused by inflammatory cytokines. When hepcidin is made to increase five-fold for a period of 365 days, the model predicts a loss of plasma iron accompanied by decreasing levels of iron in bone marrow, red blood cells (causing the anemia) and the rest of the body, while there is an increase in duodenal iron concentration (due to the inhibition of duodenal ferroportin) and transient increases in spleen and liver iron. This is in qualitative agreement with what is observed in this disease. Thus these results confirm that the action of hepcidin alone is sufficient to explain the mouse iron distribution in low iron conditions, both dietary deficiency and lowered hepcidin synthesis (e.g. as a consequence of inflammation).

In terms of the failure of the model to match the data of iron-rich diet, we should first consider whether this may be caused by the assumptions adopted in this model. One of these assumptions is the lumping the iron content of several organs into one single compartment (rest of body). Another assumption is that in each organ there is only one pool of iron (two if we consider the radioactive vs. non-radioactive forms) and there is no distinction between heme-bound iron, ferritin-bound iron and the labile iron pool. The model also does not consider the effect of the intracellular regulation that happens through transcriptional and post-transcriptional regulation (iron response elements and iron response proteins). Of all these assumptions, perhaps the one that is likely to have the strongest impact is the lack of several distinct pools of iron. In particular the simulation under-estimation of iron accumulation in the liver may be impacted by lack of ferritin and heme pools of iron; these forms of iron are not readily available to be exported out of the cell and thus could provide a higher capacity to retain iron in the liver, which the present model lacks. However, this is unlikely to be the full explanation, since the elevated hepcidin in the simulation already limits the export of iron from the liver. Additionally, the comparison of parameters between the adequate and rich diet indicated that the intake rate of iron into the liver should increase with the increase of iron in the diet. Thus it is perhaps more important to look at the details of iron import to the liver under heavy iron loads, which may be tied up with the IRE/IRP regulation in hepatocytes. Since the model is not able to appropriately explain the rich diet, specifically the high iron accumulation in the liver, we could not simulate hemochromatosis which is mainly characterized by the liver iron overload.

The failure to reproduce the red blood cell content in the iron-rich diet is more likely to be related with regulation of erythropoiesis. What would be required in the simulation to match the data would be a lower rate of iron import into the red blood cells under these conditions as predicted by the parameter comparison analysis between the models fit to the adequate and iron-rich diets. We note that in the iron-deficient diet the fit of RBC iron is much better, though it slightly under-estimates the iron content, which would go in the direction that the rate of import of iron into bone marrow is inversely proportional to the body iron.

It is only because of the rich data set of Schümann *et al.* [23] that this analysis was possible. Previously Lopes *et al.* had already produced a model based on these data [22], however since that was limited to steady state analysis it only provided three snapshots of the iron fluxes in the three different conditions and did not consider the role of hepcidin explicitly. In order to consider hepcidin it was also necessary to include the non-radioactive iron in the model. Since neither hepcidin nor non-radioactive iron were measured here, the model is less well determined than one would desire, and required adoption of a few assumptions (*e.g.* that total iron is close to 2 mg, and that the lifetime of mouse red blood cells is around 40 days).

Despite the strong effect that hepcidin has on iron absorption by enterocytes and mobilization from liver and macrophages, the present results suggest that it is not able, alone, to completely explain the iron distribution in the body. The present quantitative analysis suggests that other factors must also play a role in this process in conditions of high iron status.

This analysis of the model failures is exactly what makes the present model useful, even if in its present form it is not very predictive for diets different than the adequate. The model points to what else is important and thus it is *useful* using Box′s criterion. Its failings suggest how to improve it and gain a better understanding of iron regulation. Eventually such a model, to succeed in summarizing all that is known about this physiological process and become truly predictive, will need to include details of intracellular iron regulation – indeed we plan to follow that course and include some of those details (*e.g.* [19]) in a future multiscale model.

## Declarations

### Ethics approval

The animal data used here was previously published by other authors [23].

### Availability of data and materials

The data sets supporting the conclusions of this article are included within the article and its additional files.

### Authors′ contributions

PM conceived the study. JHP, GD, HS and PM created the model and analyzed the data. JHP and PM wrote the manuscript. All authors read the manuscript and approved its final version.

## Acknowledgements

We thank Karin Finberg and Reinhard Laubenbacher for discussions about this work. GD and HS worked on this project under the NSF Research Experience for Undergraduates project *Modeling and Simulation in Systems Biology* (DMS-1460967).

